# CCSN: Single Cell RNA Sequencing Data Analysis by Conditional Cell-specific Network

**DOI:** 10.1101/2020.01.25.919829

**Authors:** Lin Li, Hao Dai, Zhaoyuan Fang, Luonan Chen

## Abstract

The rapid advancement of single cell technologies has shed new light on the complex mechanisms of cellular heterogeneity. However, compared with bulk RNA sequencing (RNA-seq), single-cell RNA-seq (scRNA-seq) suffers from higher noise and lower coverage, which brings new computational difficulties. Based on statistical independence, cell-specific network (CSN) is able to quantify the overall associations between genes for each cell, yet suffering from a problem of overestimation related to indirect effects. To overcome this problem, we propose the “conditional cell-specific network” (CCSN) method, which can measure the direct associations between genes by eliminating the indirect associations. CCSN can be used for cell clustering and dimension reduction on a network basis of single cells. Intuitively, each CCSN can be viewed as the transformation from less “reliable” gene expression to more “reliable” gene-gene associations in a cell. Based on CCSN, we further design network flow entropy (NFE) to estimate the differentiation potency of a single cell. A number of scRNA-seq datasets were used to demonstrate the advantages of our approach: (1) one direct association network for one cell; (2) most existing scRNA-seq methods designed for gene expression matrices are also applicable to CCSN-transformed degree matrices; (3) CCSN-based NFE helps resolving the direction of differentiation trajectories by quantifying the potency of each cell. CCSN is publicly available at http://sysbio.sibcb.ac.cn/cb/chenlab/soft/CCSN.zip.

## Introduction

With the development of high-throughput single-cell RNA sequencing (scRNA-seq), novel cell populations in complex tissues [1-5] can be identified and the differentiation trajectory of cell states [6-8] can be obtained, which opens a new way to understand the heterogeneity and transition of cells [9-11]. However, compared to traditional bulk RNA-seq data, the prevalence of high technical noise and dropout events is a major problem in scRNA-seq [12-17], which raises substantial challenges for data analysis. Many computational methods were proposed to improve the identification of new cell types [18-21]. Meanwhile, imputation is an effective strategy to transform the dropouts to the substituted values [22-26]. However, most of these methods mainly analyze mRNA expression/concentrations, while the information of gene-gene interactions (or their network) is ignored.

Recently, a network-based method, cell-specific network (CSN), was proposed to perform network analysis for scRNA-seq data [27], which elegantly infers a network for each cell and successfully transforms the noisy and “unreliable” gene expression data to the more “reliable” gene association data. The network degree matrix (NDM) derived from CSN can be further applied in downstream single cell analyses, which performs better than traditional expression-based methods in terms of robustness and accuracy. CSN is able to identify the dependency between two genes from single-cell data based on statistical independence. However, CSN suffers from a problem of overestimation on gene-gene associations, which include both direct and indirect associations due to interactive effects from other genes in a network. In other words, a gene pair without direct association can be falsely identified to have a link just because they both have true associations with some other genes. Thus, the gene-gene network of a cell constructed by CSN may be much denser than the real molecular network in this cell, in particular when there are many complex associations among genes.

To overcome this shortcoming of CSN, we introduce a novel computational method to construct a conditional cell-specific network (CCSN) from scRNA-seq data. Specifically, CCSN identifies direct associations between genes by filtering out indirect associations in the gene-gene network based on conditional independence. Thus, CCSN can transform the original gene expression data of each cell to the direct and robust gene-gene association data (or network data) of the same cell. In this paper, we first demonstrate that the transformed gene-gene association data not only are fully compatible with traditional analyses such as dimension reduction and clustering, but also enable us to delineate the cell-specific network topology and its dynamics along developmental trajectories. Then, by defining the network flow entropy (NFE) on the gene-gene association data of each cell based on CCSN, we estimate the differentiation potency of individual cells. We show that NFE can illustrate the lineage dynamics of cell differentiation by quantifying the differentiation potency of cells, which is also one of the most challenging tasks in developmental biology.

## Methods

Assuming that *x* and *y* are two random variables, and *z* is the third random variable. If *x* and *y* are independent, then

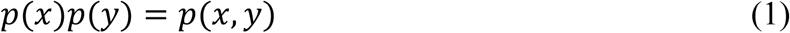

where *p*(*x, y*) is the joint probability distribution of *x* and *y*; *p*(*x*) and *p*(*y*) are the marginal probability distributions of *x* and *y.*

If *x* and *y* with the condition *z* are conditionally independent, then

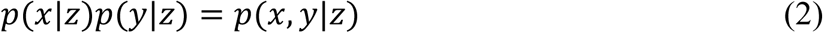

where *p*(*x, y*|*z*) is the joint probability distribution of *x* and *y* with the condition *z, p*(*x*|*z*) and *p*(*y*|*z*) are conditionally marginal probability distributions. Note that eqns. (1)-(2) are both necessary and sufficient conditions on mutual independence and conditional independence, respectively.

Here we define

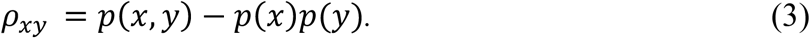

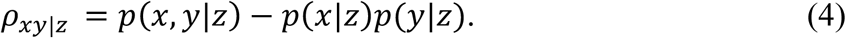

The original CSN method [27] uses *ρ*_*xy*_ to distinguish the independency and association between *x* and *y* (File S1 Note 1). However, if two independent variables *x* and *y* are both associated with a third random variable *z, ρ*_*xy*_ cannot measure the direct independency because there is an indirect association between *x* and *y.* In other words, the associations defined by CSN or eqn. (3) include both direct and indirect dependency, thus resulting in the overestimation on gene-gene associations. To overcome this problem of CSN, we develop a novel method, conditional cell-specific network (CCSN), which measures the direct gene-gene associations based on the conditional independency *ρ*_*xy*|*z*_, i.e. eqn. (4), by filtering out the indirect associations in the reconstructed network. The computational framework of CCSN is shown in **Figure 1**, and is described in the next sections.

**Figure 1.**
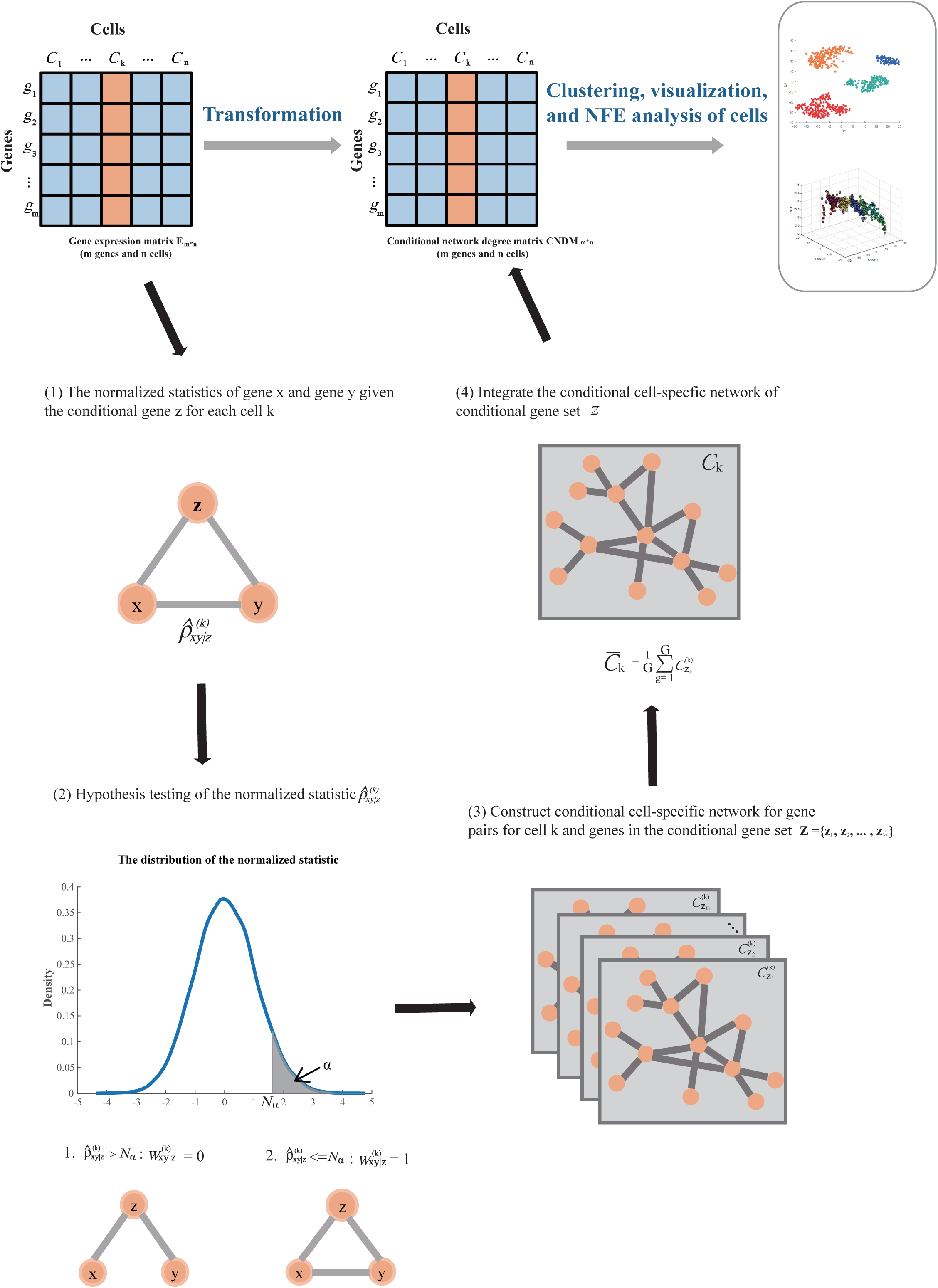
Overview of CCSN. The input data is gene expression matrix *E*_*m*\**n*_ (The orange column represents the cell *k*). (1) The normalized statistics 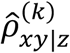 of each gene pair gene x and gene y given a conditional gene z for each cell k. 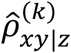 can be used to measure the direct gene-gene associations. (2) Hypothesis testing of the normalized statistic 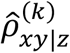. The significance level of hypothesis testing is α and 𝒩_α_ is the alpha quantile of the distribution. When 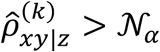, gene x and gene y are conditionally independent given the gene z in cell k, 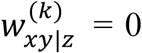, else 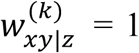. (3) Constructing conditional cell-specific network for each gene pair for cell k and for the conditional gene set 𝒵 = {*z*_1_, *z*_2_, …, *z*_*G*_}. (4) Integrating the conditional cell-specific network of conditional gene set Z. For each cell, we repeat the steps (1) – (4). Finally, we get a conditional degree matrix *CNDM* which has the same dimension as gene expression matrix *E.* The *CNDM* can be used in clustering, visualization and differentiation potency analysis.

### Probability distribution estimation

We numerically estimate the value of *ρ*_*xy*|*z*_ by making a scatter diagram based on gene expression data. Suppose there are m genes and n cells in the data. We depict the expression values of gene *x*, gene *y* and the conditional gene *z* in a three-dimensional space (Figure S1 A-G), where each dot represents one cell. First, we draw two parallel planes which are orthogonal with *z* axis near the dot *k* to represent the upper and lower bounds of the neighborhoods of *z*_*k*_. And the number of dots in the space between the two parallel planes (i.e. the neighborhood of *z*_*k*_) is 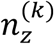 (Figure S1 D). Now we get a subspace on condition of gene z. Then, we draw other four planes near the dot *k*, where two planes are orthogonal with *x* axis and the other two planes are orthogonal with *y* axis. We can get the neighborhoods of (*x*_*k*_, *z*_*k*_), (*y*_*k*_, *z*_*k*_) and (*x*_*k*_, *y*_*k*_, *z*_*k*_) according to the intersection space of six planes (Figure S1 E-G), where the numbers of dots are 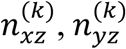 and 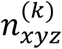, respectively. Then, we can get the estimation of probability distributions:

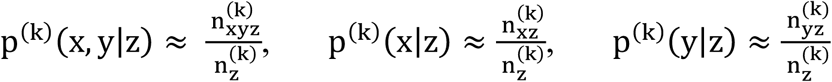

Based on eqn. (4), we construct a statistic

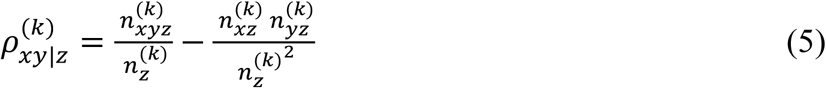

to measure the conditional independence between gene *x* and gene *y* on the condition of gene *z* in cell *k.* And when gene *x* and gene *y* given gene z are conditionally independent, the expectation 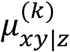 and standard deviation 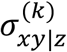 (File S1) of the statistic 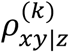 can be obtained:

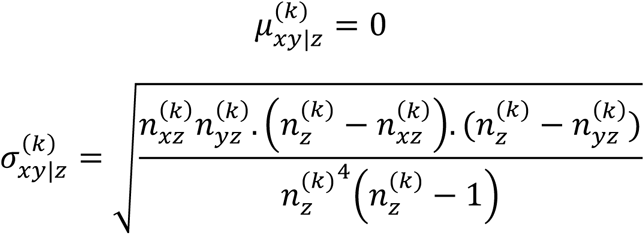

Then, we normalize the statistic as

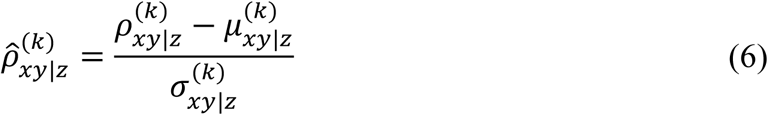

If gene *x* and *y* are conditionally independent on the condition of gene *z*, it can be proved that the normalized statistic follows the standard normal distribution (File S1 Note 1 and Figure S2), and it is less than or equal to 0 when gene *x* and *y* are conditionally independent (File S1 Note 2).

### Constructing conditional cell-specific network for each cell

To estimate the conditional independency of gene *x* and gene *y* given the conditional gene *z* in cell *k*, we use the following hypothesis test:

*H*_0_(*null hypothesis*): gene *x* and gene *y* are conditionally independent given gene *z* in cell *k.*

*H*_1_(*alternative hypothesis*): gene *x* and gene *y* are conditionally dependent given gene *z* in cell *k.*

If 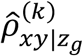, the normalized statistic, is larger than 𝒩_*α*_ (significance level α, 𝒩_*α*_ is the alpha quantile of the standard normal distribution), the null hypothesis will be rejected and then 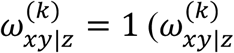 is the edge weight of genes *x* and *y* on condition of gene *z*).

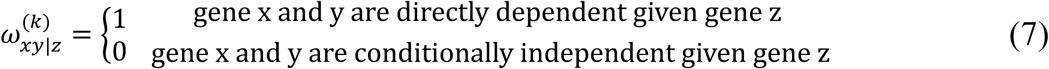

All gene pairs can be tested if they are conditionally independent given gene z in cell k. And the conditional cell-specific network (CCSN) 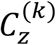 given conditional gene *z* is obtained for cell *k.*

Then, to estimate the direct association between a pair of genes in a cell, theoretically we should use all the remaining m-2 genes as conditional genes, which is computationally intensive. Suppose there are m genes in our analysis, then m*(m-1)/2 gene pairs should be tested. Fortunately, a molecular network is generally sparse, which means that a pair of genes (i.e. genes x and y) are expected to have a very small number of commonly interactive genes (as conditional genes z). In other words, numerically we can use a small number of conditional genes to identify the direct association between a pair of genes in a cell, which can significantly reduce the computational cost (File S1 Note 3, Table S1). For each gene pair in a cell, we choose G (1≤G≤m-2) genes as the conditional genes to test if the gene pair is conditionally independent or not. Generally, the conditional genes may be the key regulatory genes in a biological process, such as transcription factors and kinases. From a network viewpoint, these genes are usually hub genes in the gene-gene network, and the network degrees of these genes would be higher.

Practically, the conditional genes could be obtained from many available methods, such as highly expressed genes, highly variable genes, key transcription factor genes, or the hub genes in the CSN, and so on. For the CCSN method, the conditional gene sets were defined by CSN. The following two steps were used to obtain the conditional genes although other appropriate schemes can also be used:

1. For a given cell, we first construct a CSN without the consideration of conditional genes, where the edge between gene x and gene y in cell k is determined by the following hypothesis test:

*H*_*0*_(*null hypothesis*): gene x and gene y are independent in cell k.

*H*_*1*_*(alternative hypothesis):* gene x and gene y are dependent in cell k.

The statistic *ρ*_*xy*_ can be used to measure the independency of genes x and y (File S1 Note1). If *ρ*_*xy*_ is larger than a significant level, we will reject the null hypothesis and edge_xy_ (k) = 1, otherwise edge_xy_ (k) = 0.

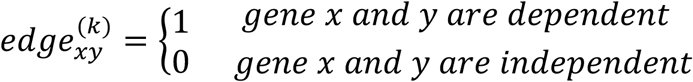

Then we use 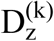 to measure the importance of conditional gene z in cell k:

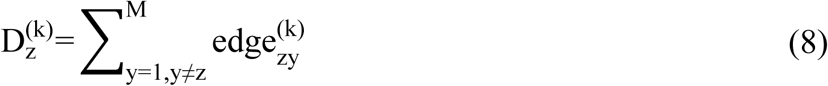

Eqn. (8) means that if a gene is connected to more other genes, this gene is more important.

2. For a given cell k, we choose the top *G* (*G* ≥ 1) largest ‘importance’ genes as the conditional genes.

We assume that the conditional gene set is {*z*_*g*_, g = 1,2,3, …, G}, and the conditional cell-specific network (CCSN) 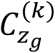 is obtained for cell *k* given conditional gene *z*_*g*_. The CCSNs of the cell k on the condition of gene set {*z*_*g*_, g = 1,2,3, …, G} are 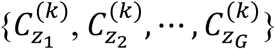. Then, we use

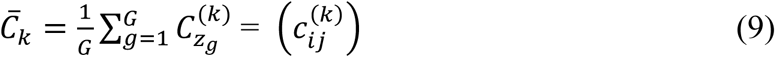

to represent the degrees of gene-gene interaction network of cell *k*, where 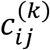 for *i, j* = 1, …, *m* is the (i,j) element of the matrix 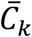.

For scRNA-seq data with all *n* cells, we can construct *n* CCSNs, which can be used for further dimension reduction and clustering. In other words, instead of the originally measured gene expression data with *n* cells, we use the *n* transformed CCSNs for further analysis. In addition, each CCSN is a network for a cell, which can be used for network analysis (gene regulations and network biomarkers) on the basis of a single cell.

### Network degree matrix from CCSN

CCSNs could be used for various biological studies by exploiting the gene-gene conditional association network from a network viewpoint. We transform eqn. (9) to a conditional network degree vector based on the following transformation

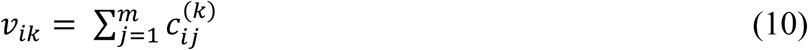

Then, for 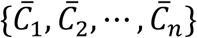 an *m*\**n* matrix CNDM is obtained.

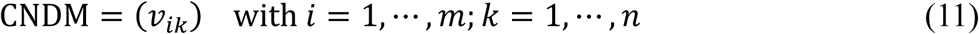

The matrix has the same dimension with the gene expression matrix (GEM), i.e. GEM= (*x*_*ik*_) with *i* = 1, …, *m*; *k* = 1, …, *n*, but CNDM can reflect the gene-gene direct association in terms of interaction degrees. Moreover, this CNDM matrix after normalization could be further analyzed by most traditional scRNA-seq methods for dimension reduction and clustering analysis. The input/output settings as well as application fields of our CCSN method are listed in File S1 Note 4.

### Network analysis of CCSN

The relationship between gene pairs can be obtained by CCSN at a single cell level. CCSN also provides a new way to build gene-gene interaction network for each cell. And the CNDM derived from CCSN can be further used in dimension reduction, clustering and network flow entropy analysis by many existing methods.

#### Dimension reduction

We used principal component analysis (PCA) [28] and t-distributed stochastic neighbor embedding (t-SNE) [29] which respectively represent linear and nonlinear methods, to perform dimension reduction on public scRNA-seq datasets with known cell types.

#### Clustering

To validate the good performance of CCSN in clustering analysis, several traditional clustering methods such as K-means, Hierarchical cluster analysis, and K-medoids were applied to clustering analysis. Furthermore, state-of-the-art scRNA-seq data clustering methods such as SC3, SIMLR and Seurat [20, 30, 31] were also used for comparison.

#### Network flow entropy analysis

Quantifying the differentiation potency of a single cell is one of the important tasks in scRNA-seq studies [15, 32, 33]. A recent study developed SCENT [34], which uses protein-protein interaction (PPI) network and gene expression data as input to obtain the potency of cells. However, SCENT depends on the PPI network, which may ignore many important relationships between genes in specific cells. In this paper, we developed network flow entropy (NFE) to estimate the differentiation potency of a cell from its CSN or CCSN network, which is constructed for each cell. The normalized gene expression profile and CSN/CCSN is used when we compute the network flow entropy.

Estimating NFE requires a background network, which could be provided by CSN or CCSN. Based on CSN or CCSN, we could know whether or not there is an edge between gene i and gene j. We assume that the weight of an edge between gene *i* and gene *j, p*_*ij*_ is proportional to the normalized expression levels of gene *i* and gene *j*, that is *p*_*ij*_ ∝ *x*_*i*_*x*_*j*_ with 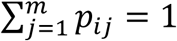. These weights are interpreted as interaction probabilities. Then, we normalize the weighted network as a stochastic matrix, *P=*(*p*_*ij*_) with

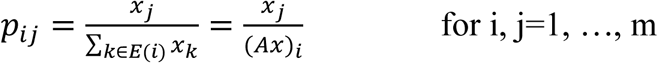

where *E*(*i*) contains the neighbours of gene *i*, and *A* is the CSN or CCSN (*A*_*ij*_ = 1 if *i* and *j* are connected, otherwise *A*_*ij*_ = 0).

And then, we define the NFE as:

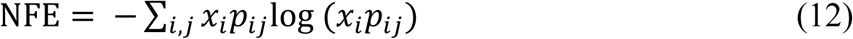

where *x*_*i*_ is the normalized gene expression of gene *i.* From the definition, NFE is clearly different from network entropy.

### Data availability

Twelve scRNA-seq datasets and one bulk RNA-seq dataset [15, 35-41] were used to validate our CCSN method. The numbers of cells in these datasets range from 100 to 20,000. Table S2 gives a brief introduction of these datasets.

## Results

### Visualization and clustering of scRNA-seq datasets with CNDM

Characterizing the cell heterogeneity is one of the important tasks for scRNA-seq data analysis. To test whether CCSN-transformed network data can help segregate cell types, we performed dimension reduction and clustering on the CNDMs of gold-standard scRNA-seq datasets, using algorithms widely employed in scRNA-seq studies. The numbers of conditional genes used in CCSN construction were listed in Table S2.

For visualizing the structure of these datasets in a two-dimensional space, we used the representative linear and nonlinear dimension reduction methods, principle component analysis (PCA) [42] and t-distributed stochastic neighbor embedding (t-SNE) [29], respectively. As shown in **Figure 2** and Figure S3, CNDMs can separate different cell types clearly in the low-dimensional space by both PCA and t-SNE. Notably, they generally perform even better than GEM (Figure 2, Figure S3). Hence, the network data of CNDMs contain sufficient information for separating cell types in scRNA-seq datasets.

**Figure 2.**
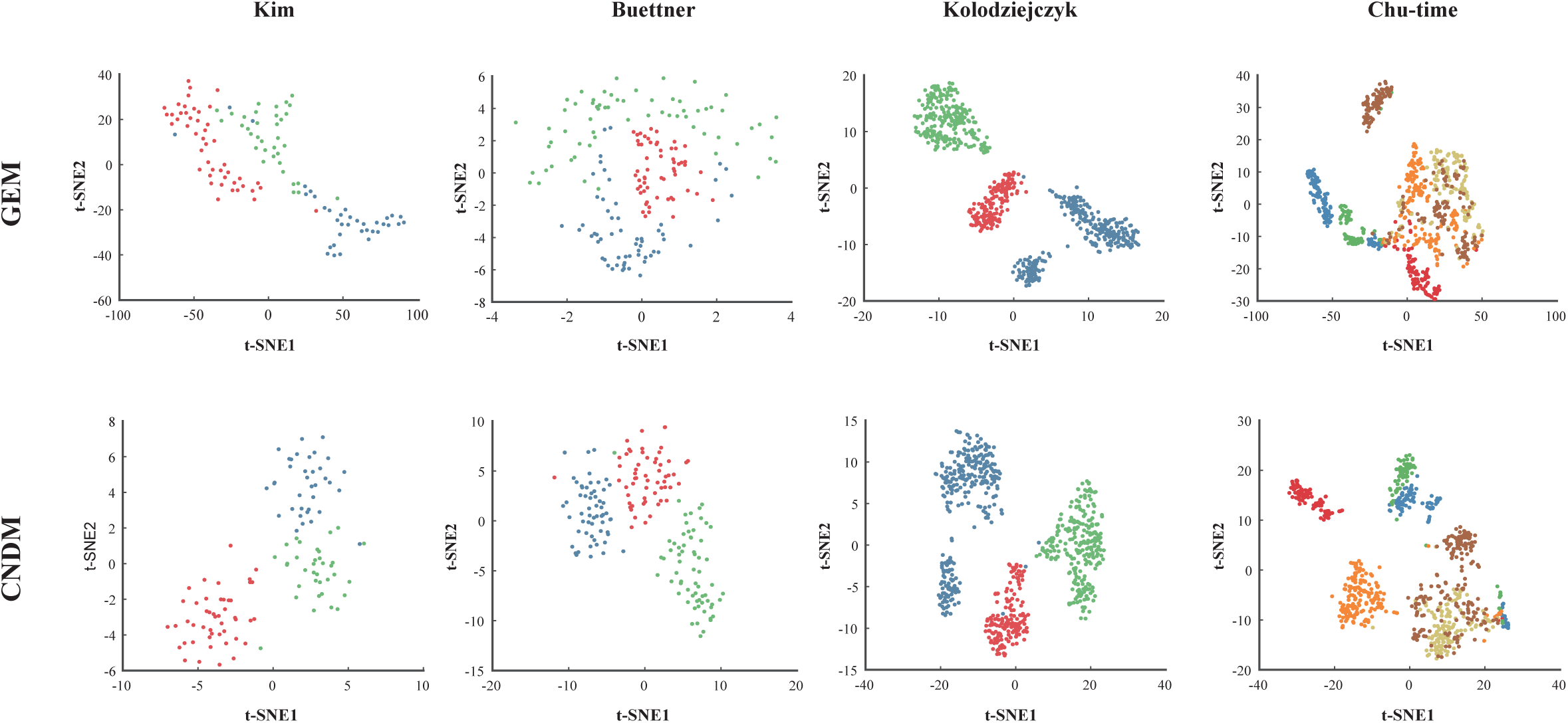
CNDM for visualization of scRNA-seq data. The datasets are dimensionally reduced by t-SNE and cell types are encoded by different colors.

To quantitatively evaluate the power of CNDMs in cell type identification, we performed clustering on CNDMs and computed the adjusted random index (ARI) for each dataset based on the background truth (File S1 Note 5). As shown in **Table 1** and Figure S4, CNDMs perform obviously better than GEM on all datasets, either without or with dimension reduction with t-SNE. These provide a strong support of the notion that the CCSN-transformed network data are highly informative for characterizing single cell populations. Interestingly, when further compared to NDM, CNDMs also show a good performance (**Table 2** and Figure S5).

**Table 1.** The comparison of CNDM and GEM in clustering of scRNA-seq data.

|  |  | Buettner | Kolodziejczyk | Grokce | Chu-time | Chu-type | Kim |
| --- | --- | --- | --- | --- | --- | --- | --- |
| K-means | GEM | 0.29 | 0.54 | 0.42 | 0.17 | 0.22 | 0.20 |
|  | **CNDM** | **0.87** | **0.85** | **0.75** | **0.45** | **0.57** | **0.81** |
| Hierarchical | GEM | 0.32 | 0.49 | 0.47 | 0.22 | 0.22 | 0.12 |
|  | **CNDM** | **0.73** | **0.65** | **0.92** | **0.47** | **0.61** | **0.77** |
| K-means (t-SNE) | GEM | 0.41 | 0.87 | 0.43 | 0.33 | 0.55 | 0.53 |
|  | **CNDM** | **0.95** | **0.91** | 0.36 | 0.56 | **0.70** | **0.93** |
| Hierarchical (t-SNE) | GEM | 0.55 | 0.99 | 0.50 | 0.39 | 0.67 | 0.73 |
|  | **CNDM** | **0.95** | 0.99 | 0.39 | **0.61** | **0.80** | **0.95** |
| K-medoids | GEM | 0.23 | 0.29 | 0.40 | 0.33 | 0.33 | 0.79 |
|  | **CNDM** | **0.53** | **0.63** | **0.81** | 0.17 | **0.38** | **0.61** |
| SC3 | GEM | 0.89 | 1 | 0.56 | 0.66 | 0.78 | 0.89 |
|  | **CNDM** | **0.98** | 0.72 | **0.72** | 0.63 | **0.98** | **0.96** |
| SIMLR | GEM | 0.89 | 0.49 | 0.43 | 0.30 | 0.48 | 0.38 |
|  | **CNDM** | 0.63 | **0.52** | **0.85** | **0.58** | **0.54** | **0.95** |
| Seurat | GEM | 0.67 | 0.43 | 0.35 | 0.52 | 0.52 | 0.41 |
|  | **CNDM** | **0.90** | **0.56** | 0.32 | **0.56** | **0.69** | **0.84** |

*Note:* The performance of clustering is evaluated by adjusted random index (ARI). Hierarchical (t-SNE) and k-means (t-SNE) represent that the clustering analysis is performed after dimension-reduction by t-SNE

**Table 2.** The comparison of CNDM with NDM in clustering analysis.

|  |  | Buettner | Kim | Wang | Grokce | Tabula Muris<br>(Aorta) | Tabula Muris<br>(Limb Muscle) |
| --- | --- | --- | --- | --- | --- | --- | --- |
| K-means | NDM | 0.50 | 0.50 | 0.30 | 0.79 | 0.21 | 0.58 |
|  | **CNDM** | **0.87** | **0.81** | **0.45** | **0.75** | **0.63** | **0.66** |
| Hierarchical | NDM | 0.69 | 0.59 | 0.38 | 0.95 | 0.12 | 0.65 |
|  | **CNDM** | **0.73** | **0.77** | **0.45** | **0.92** | **0.75** | **0.76** |
| K-means (t-SNE) | NDM | 0.83 | 0.84 | 0.61 | 0.38 | 0.46 | 0.62 |
|  | **CNDM** | **0.95** | **0.93** | **0.67** | 0.36 | **0.61** | **0.65** |
| Hierarchical (t-SNE) | NDM | 0.89 | **0.98** | 0.58 | 0.47 | 0.50 | 0.66 |
|  | **CNDM** | **0.95** | **0.95** | **0.72** | 0.39 | **0.50** | 0.66 |
| K-medoids | NDM | 0.26 | 0.49 | 0.31 | 0.60 | 0.35 | 0.14 |
|  | **CNDM** | **0.53** | **0.61** | 0.21 | **0.81** | **0.53** | **0.39** |
| SC3 | NDM | 0.67 | 1 | 0.70 | 0.45 | 0.29 | 0.66 |
|  | **CNDM** | **0.98** | 0.96 | **0.86** | **0.72** | **0.73** | **0.76** |
| SIMLR | NDM | 0.64 | 0.75 | 0.29 | 0.74 | 0.40 | 0.60 |
|  | **CNDM** | 0.63 | **0.95** | **0.60** | **0.85** | **0.70** | **0.71** |
| Seurat | NDM | 0.82 | 0.97 | 0.59 | 0.44 | 0.45 | 0.66 |
|  | **CNDM** | **0.90** | 0.84 | 0.59 | 0.32 | **0.76** | **0.75** |

*Note:* The performance of clustering is evaluated by adjusted random index (ARI). Hierarchical (t-SNE) and k-means (t-SNE) represent that the clustering analysis is performed after dimension-reduction by t-SNE

To further evaluate CCSN for larger datasets, the Tabula Muris droplet1 dataset [41] comprising more than 20,000 cells from three tissues (bladder, trachea, and spleen) were tested. The Seurat package was used to perform dimension reduction and clustering analysis on the CNDM [31]. The cells are clearly segregated into three dominant groups on the t-SNE map, which are largely defined by their cell origins (ARI = 0.73 and Figure S6). This indicates that CCSN can be effectively extended to larger datasets in addition to the relatively small gold standard datasets benchmarked above.

### CCSN reveals network structure and dynamics on a single cell basis

In this paper, we apply CCSN to Wang dataset [39], which comes from a study of neural progenitor cells (NPCs) that differentiate into mature neurons. The dataset contains six time points over a 30-day period.

The CSN and CCSN are performed on a single cell (Day 0, RHB1742_d0) using 195 transcription factors which are differentially expressed across all the cell subpopulations and all time points. In CCSN, two genes (*HMGB1* and *SOX11*) of high coefficients of variation (CV) are chosen as the conditional genes. The results (**Figure 3**A) illustrate that the network of CCSN are much sparser than the network of CSN. There are three modules in the CCSN, while there is only one dense network in the CSN. Furthermore, three hub genes are obtained in three modules in CCSN. One of the hub genes is *ASCL1* which plays an important role in neural development [13, 43]. Thus, by removing indirect associations, CCSN can extract a more informative network structure than CSN, which could improve the characterization of key regulatory factors in individual cells.

**Figure 3.**
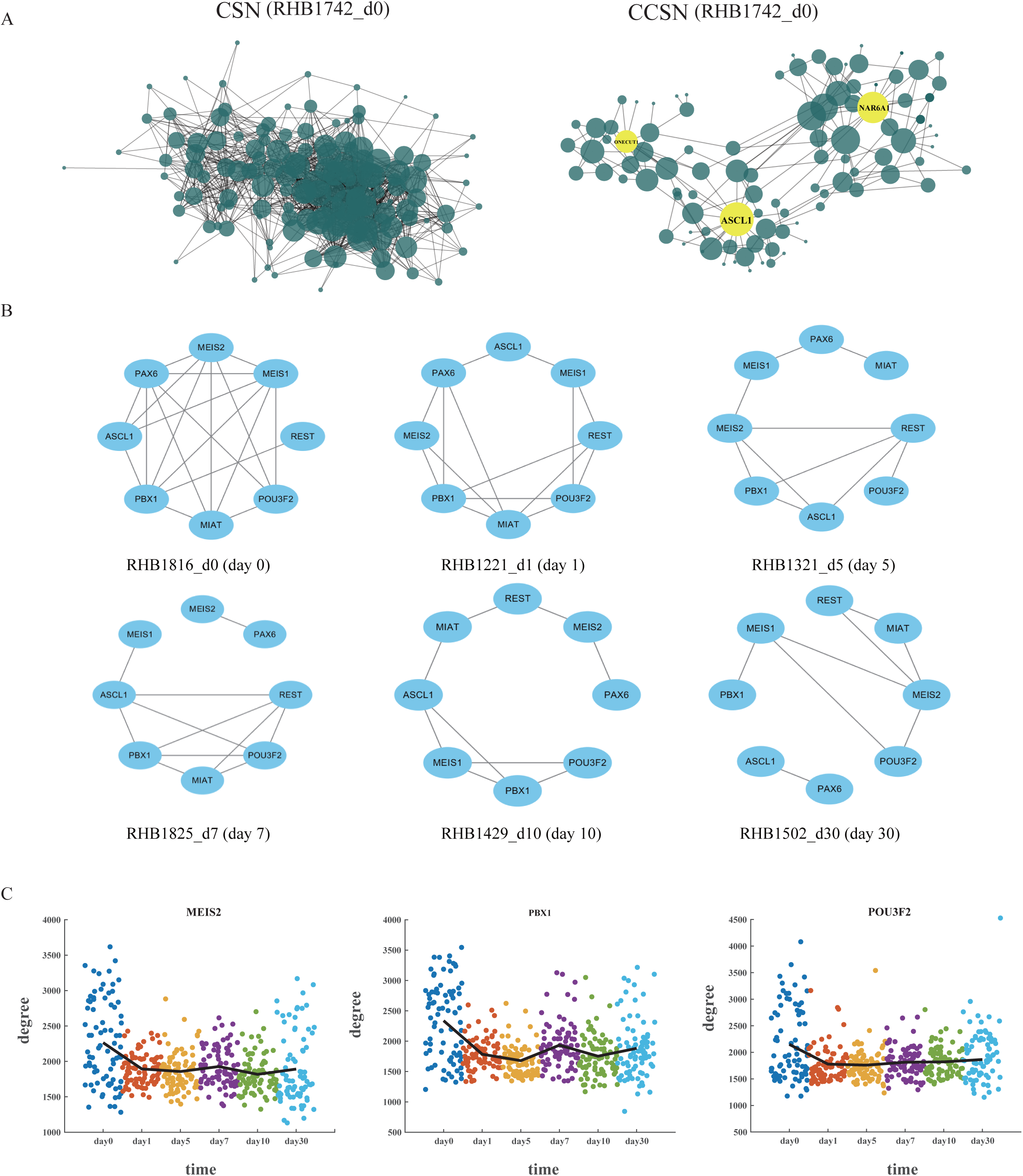
CCSN uncovers network topology and dynamics for single cells. **A**. The cell specific network (CSN) and conditional cell specific network (CCSN) of the same single cell from the Wang dataset. The same genes are used in network construction. **B**. CCSNs of 8 core genes for representative single cells. **C**. CCSN degrees of *MEIS2, PBX1* and *POU3F2* along six time points of the neuronal differentiation.

CCSN also reveals the network dynamics over the differentiation trajectory. As illustrated in Figure 3B, a core neural differentiation network composed of eight regulatory genes is dynamically modulated through the temporal progression of NPC differentiation. At day 0, the associations among these genes are the strongest, consistent with the high potency of progenitor cells. As NPC differentiates, the network becomes much sparser, suggesting more specified cell fates. In addition, when constructing CCSN from all genes, the degrees of *MEIS2, PBX1* and *POU3F2* are also larger in day 0 and quickly decreases afterwards (Figure 3C), indicating that these genes are highly connected with other genes in NPCs, consistent with their known important roles in early differentiation of neural progenitor cells [39].

Both theoretically and computationally, CCSN can also construct a gene-gene network for a single bulk RNA-seq sample, in addition to a single cell. To validate this biologically, we apply CCSN to the TCGA lung adenocarcinoma (LUAD) RNA-seq dataset. The t-SNE plot based on CNDM reveals two obvious clusters, which respectively corresponding to normal adjacent lung tissues and lung tumors (Figure S8A), supporting the effective application of CCSN to bulk RNA-seq data as well. Moreover, the EGFR pathway, a well-known oncogenic driver pathway for LUAD [44-46], is densely connected in tumor samples but not in benign tissues, as illustrated in the representative single-sample EGFR networks (Figure S8 B), and the CCSN degrees of EGF and EGFR in each normal and tumor samples (Figure S8 C). These data demonstrate that CCSN well extends to single sample bulk RNA-seq data analysis and uncovers important biological connections related to disease states.

### CCSN-based network flow entropy analysis

To quantify the differentiation state of cells, we further develop a new method “network flow entropy” (NFE) to estimate the differentiation potency of cells by exploiting the gene-gene network constructed by CCSN.

To assess the performance of NFE, we apply it to two datasets. In Wang dataset [39], there are 484 cells with 6 stages (day 0, day 1, day 5, day 7, day 10, day 30) and the CCSNs with one conditional gene are used to compute the network flow entropy. We compared NPC (at Day 0 and Day 1) with mature neurons (at Day 30) (**Figure 4**A). In Yang dataset [38], we compared the cells in day 10 with day 17 in differentiation of mouse hepatoblasts (Figure 4B) and the CSN was used to compute the network flow entropy. In both datasets, NFE assigns significantly higher scores to the progenitors than the differentiated cells (one-sided Wilcox rank sum test, p-value = 3.756e-19 in Yang dataset, p-value = 2.062e-12 on Wang dataset).

**Figure 4.**
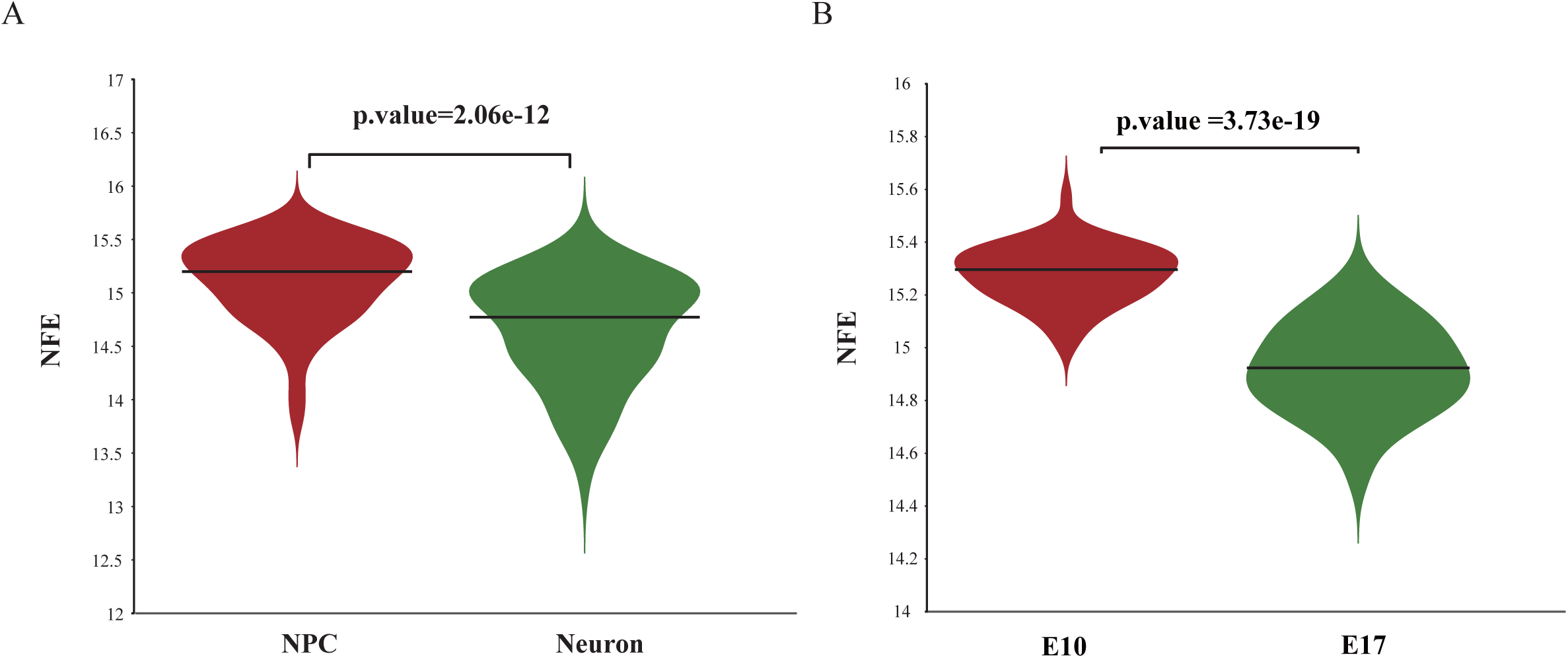
Network flow entropy analyses for differentiated cells and progenitors. **A**. Network flow entropy between NPCs (at 0 and 1 day) and mature neurons (at 30 day). **B**. Network flow entropy between cells at day 10 and day 17 during differentiation of mouse hepatoblasts. P-value is from one-sided Wilcoxon rank-sum test.

To further validate NFE, we generated a three-dimensional representation of the cell-lineage trajectory for the Wang dataset. In the time-course differentiation experiment of NPCs into neurons [39], NFE correctly predicted a gradual decrease in differentiation potency (**Figure 5**). Therefore, NFE is effectively applicable to single cell differentiation studies and highly predictive of developmental states and directions.

**Figure 5.**
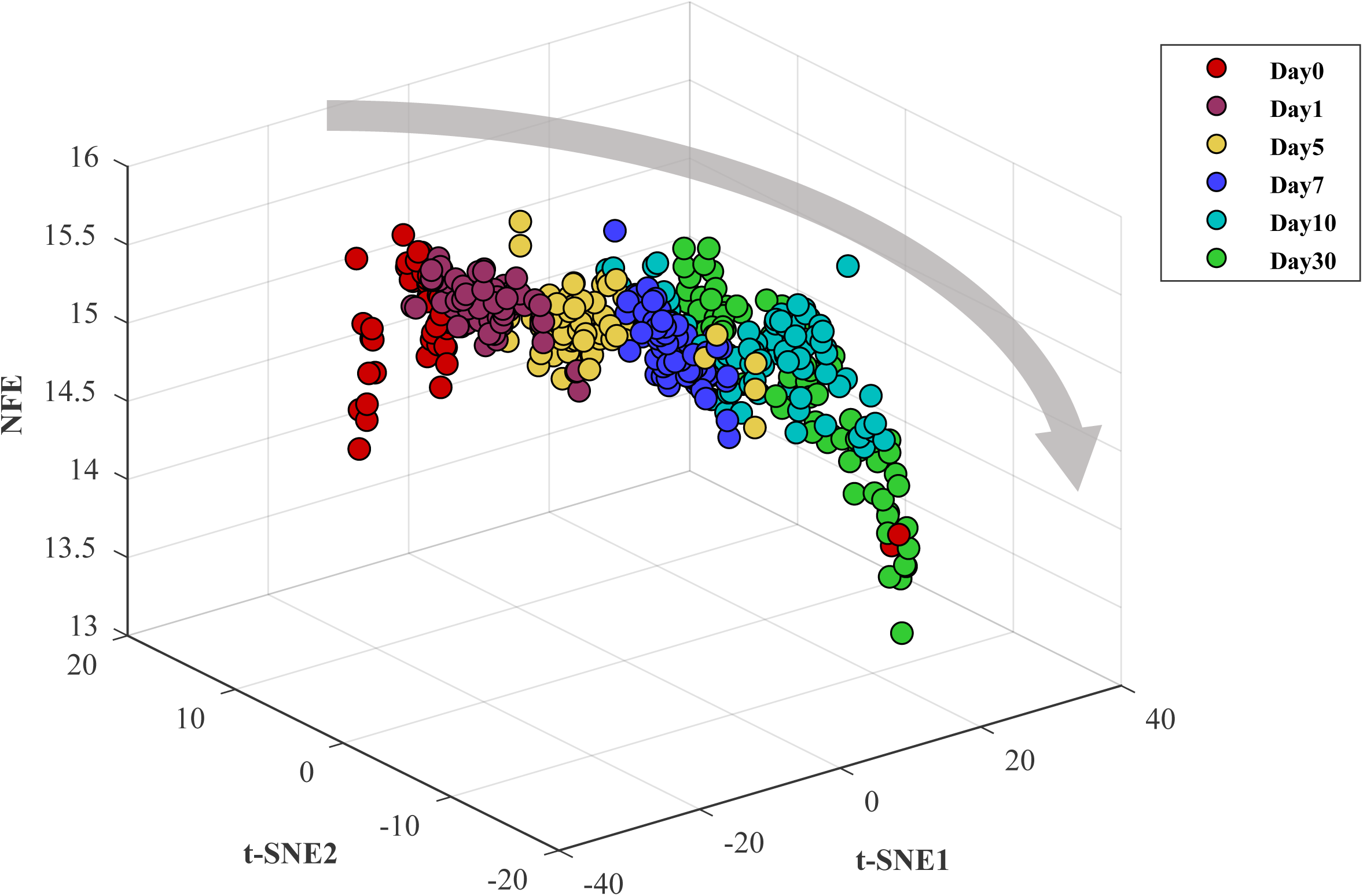
The differentiation landscape of neural progenitor cells into mature neurons. The 3-dimensional plot shows the NFE of single cells gradually decrease along the differentiation time-course of neural progenitor cells (day 0) into mature neurons (day 30). The z axis represents the NFE. The x axis and y axis are the two components of t-SNE.

## Discussion

Estimating functional gene networks from noisy single cell data has been a challenging task. Motivated by network-based data transformation, we have previously developed CSN to uncover cell-specific networks and successfully applied it to extract biologically important gene interactions. However, CSN does not distinguish direct and indirect associations and thus suffers from the so-called overestimation problem. In this study, we propose a more sophisticated approach termed CCSN, which constructs direct gene-gene associations (network) of each cell by eliminating false connections introduced by indirect effects.

CCSN can transform GEM to CNDM for downstream dimension reduction and clustering analysis. These allow us to identify cell populations, generally better than GEM in the datasets tested above. In addition, CCSN also shows good performance when compared to CSN. Moreover, we can construct one direct gene-gene association network by one cell based on CCSN. From the networks of the individual cells, we can obtain the dynamically changed networks. In Figure 3C, the CCSNs of these cells dynamically changed at different time points, and the network at day 0 shows the strongest associations. Moreover, the hub genes of the networks constructed by CCSN method may play an important role in biological processes. In Figure 3A, the hub genes of three modules in the network constructed by CCSN play a vital role in neural development. These clearly demonstrate the advantages of CCSN. Furthermore, we develop a new NFE index which can accurately estimate the differentiation potency of a single cell. And the results show that NFE performs well in distinguishing various cells of differential potency.

Nonetheless, the computational cost of CCSN generally increases by G times comparing with the original CSN due to G conditional genes. Thus, a parallel computation scheme is desired to reduce the computation time. Also, CCSN is not designed to construct the causal gene association networks, and the directions of the gene associations cannot be obtained. These could be our future research topics.

## Author Contributions

L.L. and H.D. developed the methodology. L.L. executed the experiments. Z.Y.F. helped the experiments and provided technical support. L.L., H.D., Z.Y.F. and L.N.C. wrote and revised the manuscript. L.N.C. and Z.Y.F. supervised the work and critically reviewed the paper. All authors have read and approved the final manuscript.

## Acknowledgements

This work was supported by the National Key R&D Program of China (No. 2017YFA0505500), National Natural Science Foundation of China (Nos. 31771476 and 31930022), and Shanghai Municipal Science and Technology Major Project (No. 2017SHZDZX01). We would like to acknowledge Dr. Tang Zeng for helpful discussions.

## Supplementary material

**File S1 CCSN additional implementation details**

**Figure S1** Scatter diagram of the expression values of gene x, gene y and gene z for cell k

(A) the red plot k represents the cell k and x axis, y axis and z axis represent the expression levels of gene x, gene y and gene z. gene z respectively. Gene z is set as the conditional gene. n is the number of cells in the dataset. (B) The two parallel light shadow planes 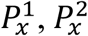, where x-axis is orthogonal with two planes. The dots are contained in the space between the two planes are the neighbors of *x*_*k*_ and the number of the dots is 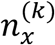. (C) The two parallel light shadow planes 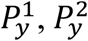, where y-axis is orthogonal with two planes. The dots are contained in the space between the two planes are the neighbors of *y*_*k*_ and the number of the dots is 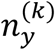. (D) The two parallel light shadow planes 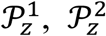, where z-axis is paralleled with the two planes. The dots contained in the space between the two planes are the neighbors of *z*_*k*_, and the number of the dots is 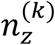. (E)The intersection of the four parallel light shadow planes 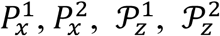 is the space which is surrounded by the green lines. The number of dots which are contained in the space is 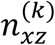. (F)The intersection of the four parallel light shadow planes 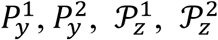 is the space which is surrounded by the green lines. The number of dots which are contained in the space is 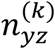. (G) The intersection of the six parallel shadow planes 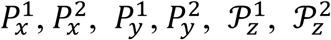 is the space which is surrounded by the green lines. The number of dots which are contained in the space is 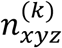.

**Figure S2** The comparison of standard normal distribution and the distribution of 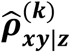

The density function is calculated by kernel density estimation based on 20,000 plots, and *n*_*x*_, *n*_*y*_, *n*_*z*_ are equal to 0.2n. The gene x and gene y are conditional independent given gene z.

**Figure S3** Performance comparison of GEM and CNDM

PCA was applied for visualization and different colors represent different cell types.

**Figure S4** The clustering performance of CNDM and GEM

K-means, hierarchical clustering algorithm (HCA) and K-medoids were used for comparison. The data which was preprocessed by t-SNE was also performed to cluster.

**Figure S5** The clustering performance of CNDM and NDM

K-means, hierarchical clustering algorithm (HCA) were used for comparison. The data which was preprocessed by t-SNE was also performed to cluster.

**Figure S6 Visualization of 23,321 cells by t-SNE**

Different colors represent different tissues.

**Figure S7 The clustering performance of CNDM with different parameters**

**Figure S8 CCSN analysis of TCGA-LUAD dataset**

A. t-SNE plots are used for visualization based on CCSN. The normal samples and tumor samples are represented by different colors. B. CCSNs of representative samples for 18 genes involved in the EGFR pathway. C. Conditional network degrees of EGF and EGGR in the normal samples and the tumor samples.

**Table S1 The running time of CCSN with different numbers of conditional genes**

**Table S2 Datasets used in this study**

