## Supplementary figures and images for "CCSN: Single Cell RNA Sequencing Data Analysis by Conditional Cell-specific Network"

### Figure S1

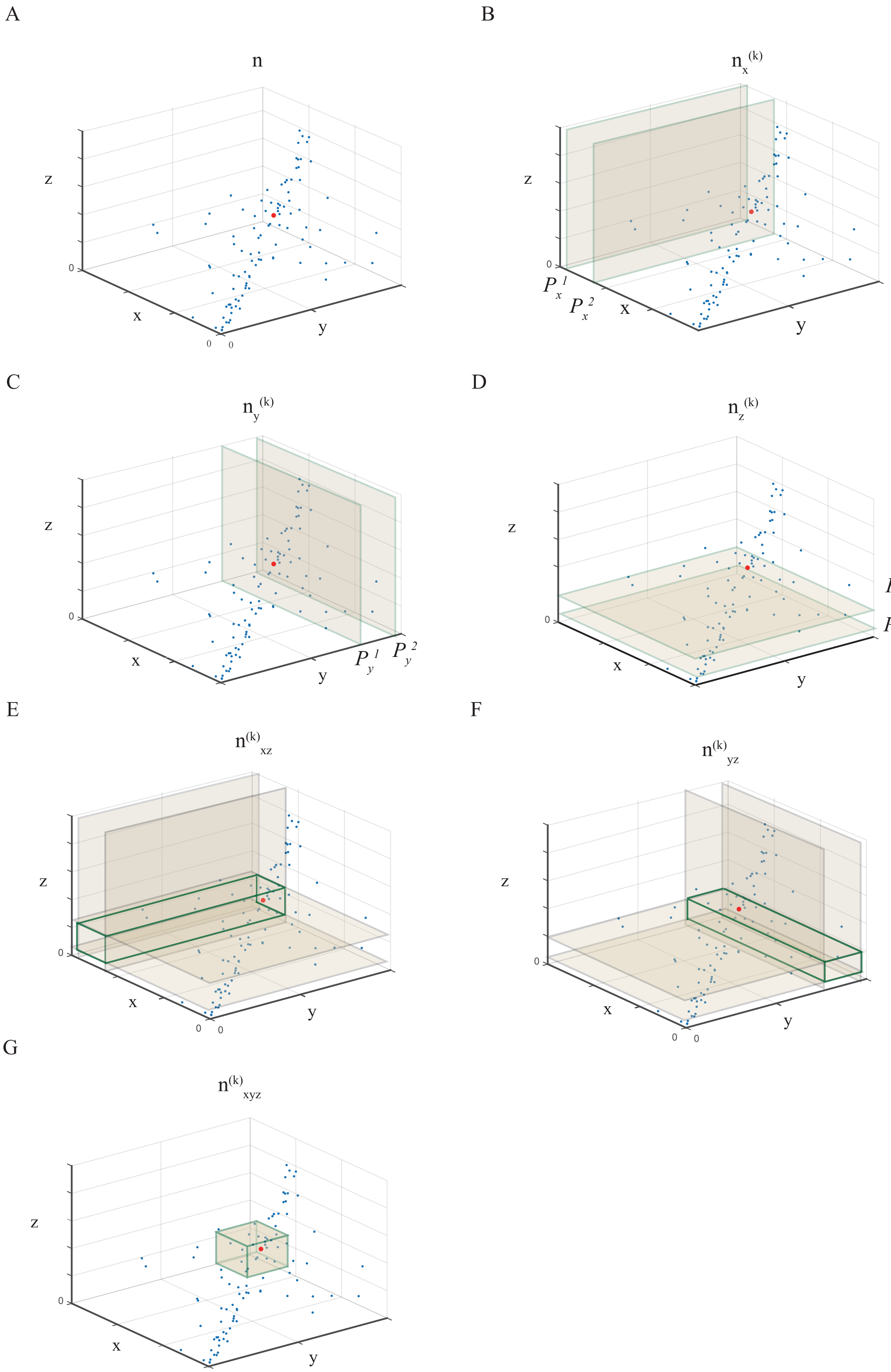

### Figure S2

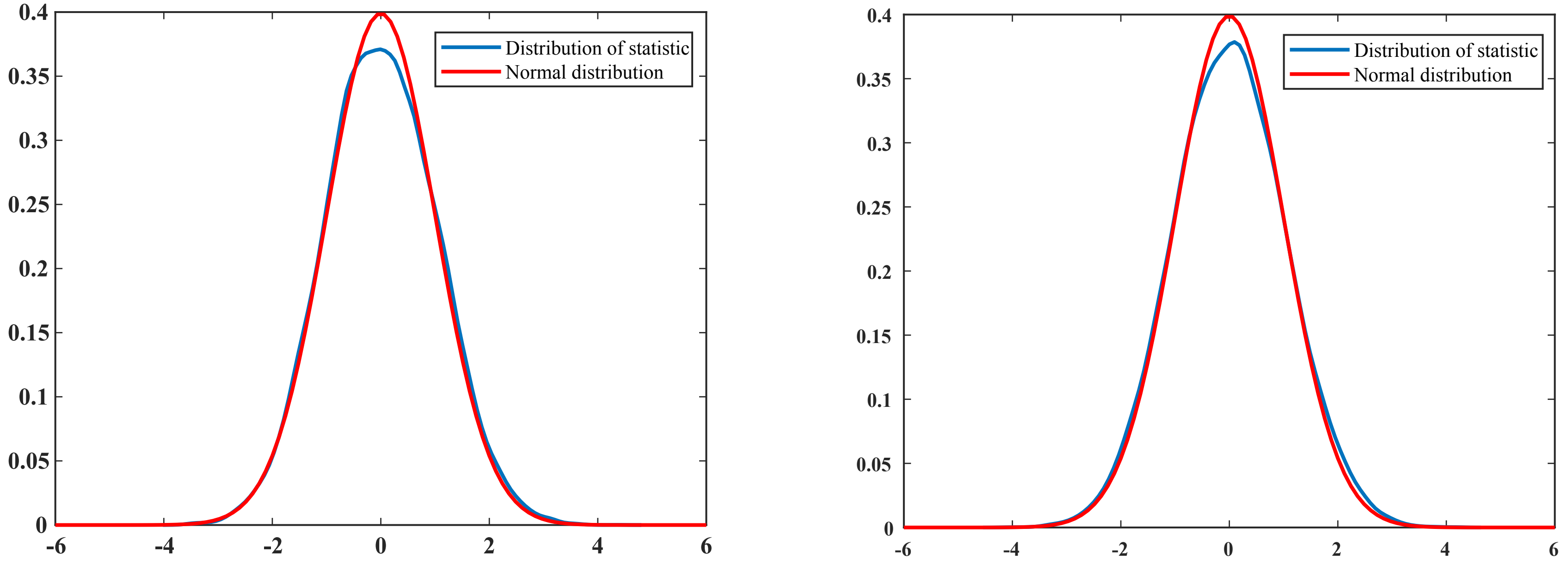

### Figure S3

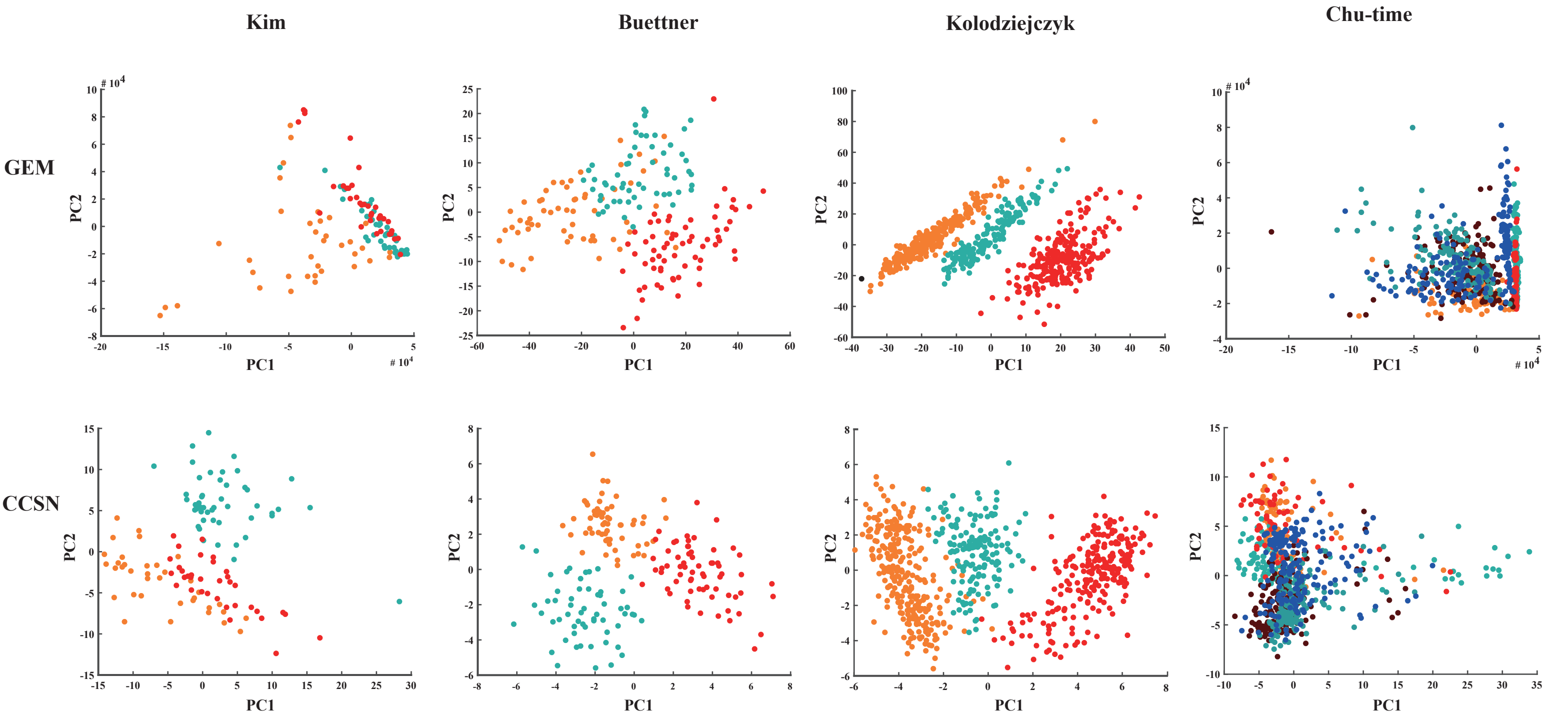

### Figure S4

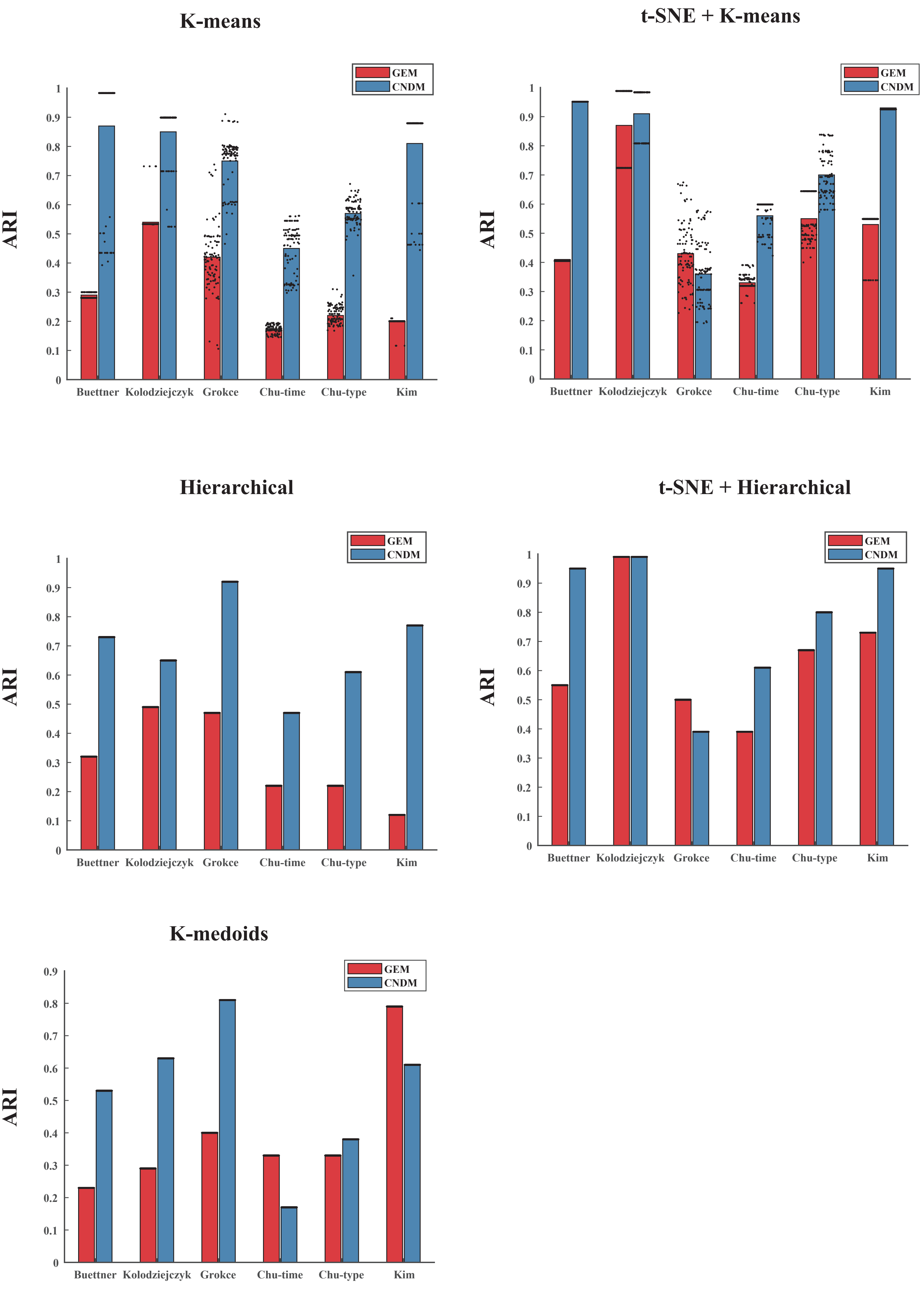

### Figure S5

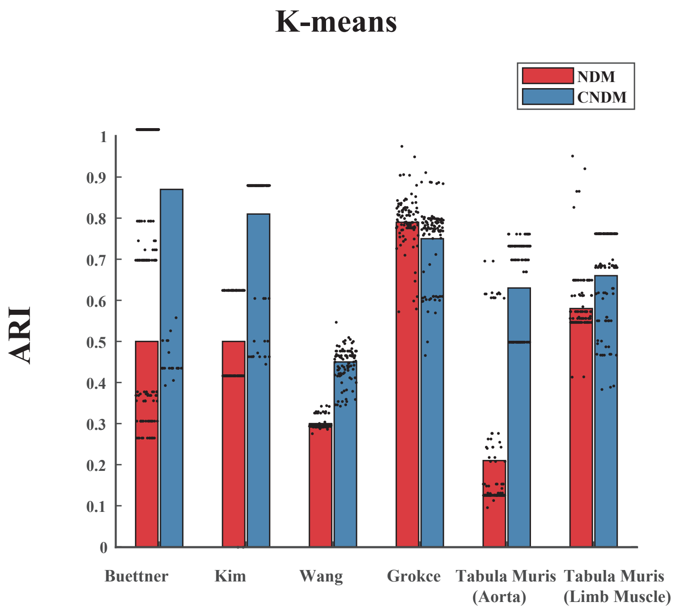

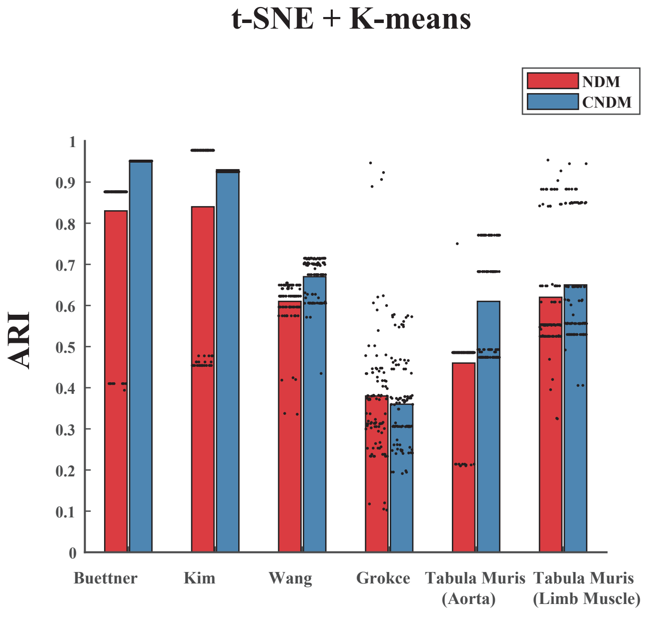

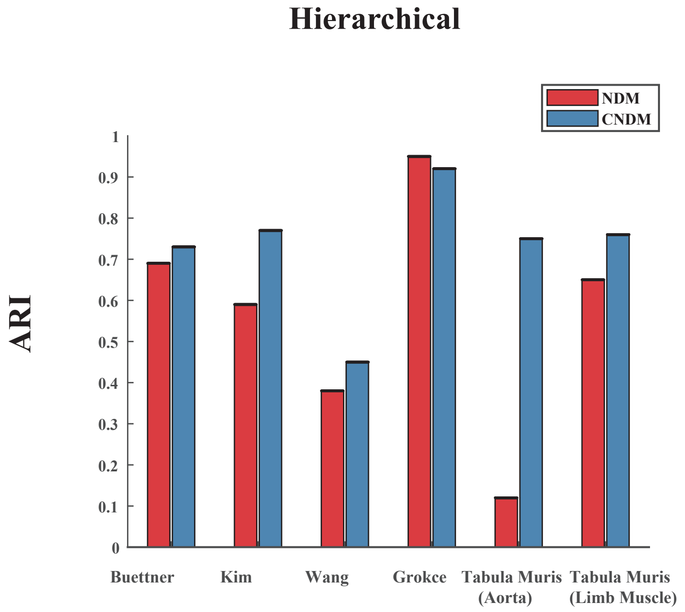

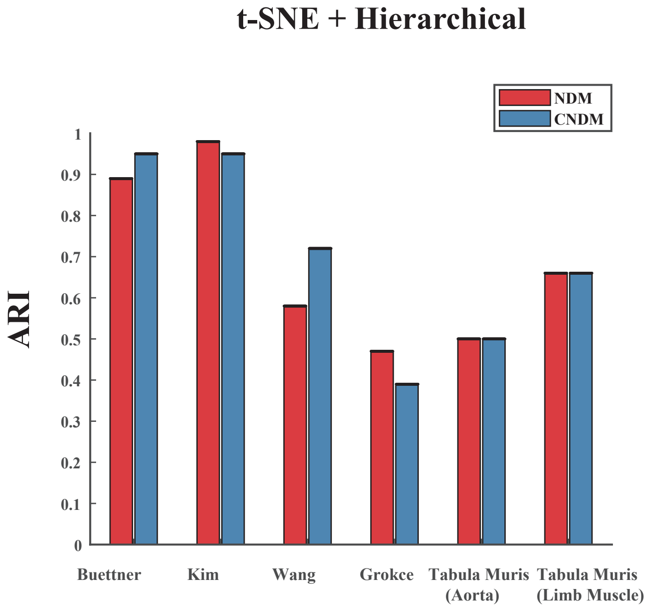

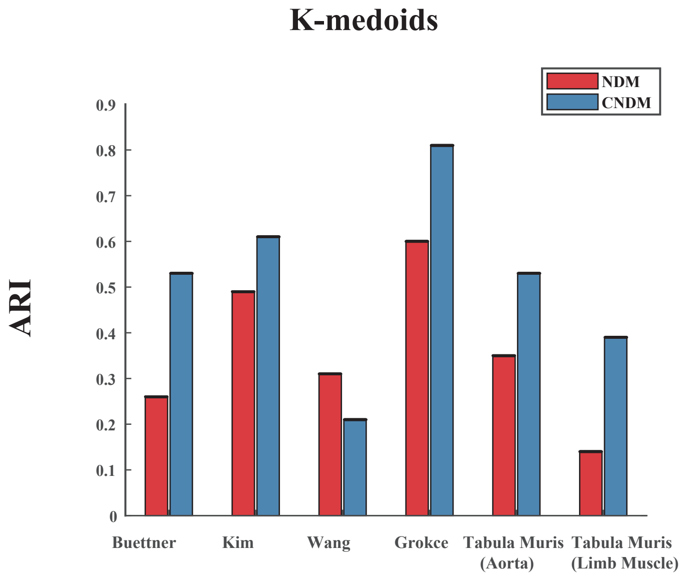

### Figure S6

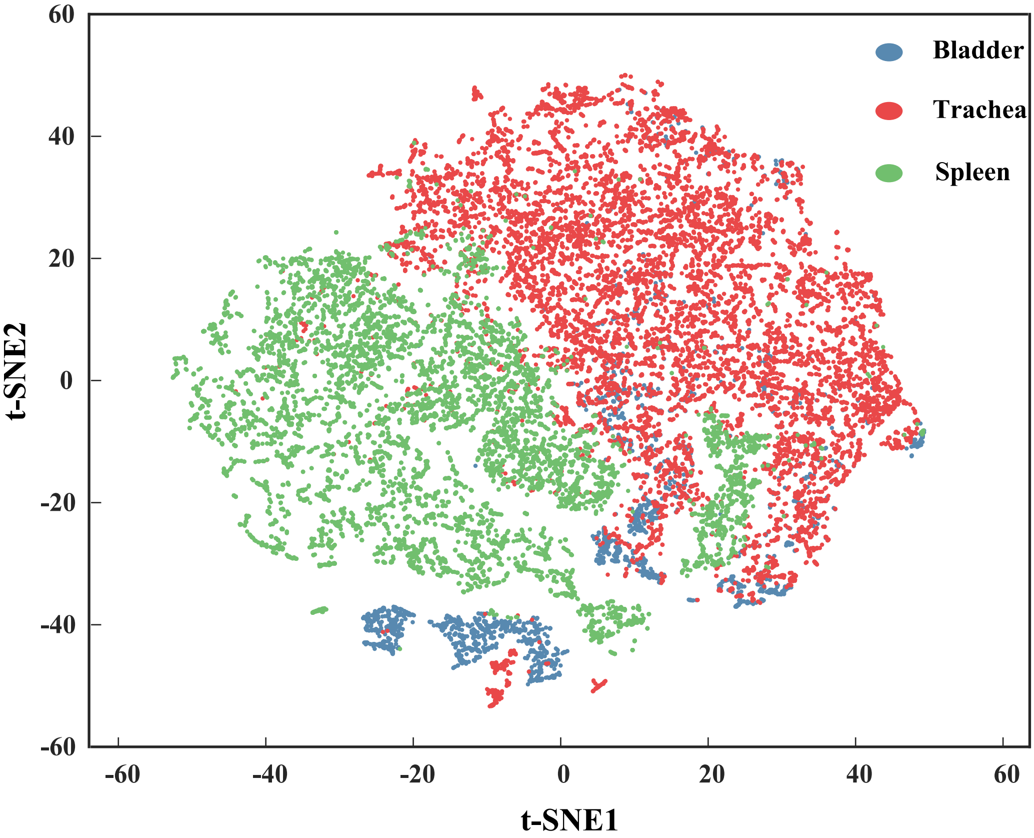

### Figure S7

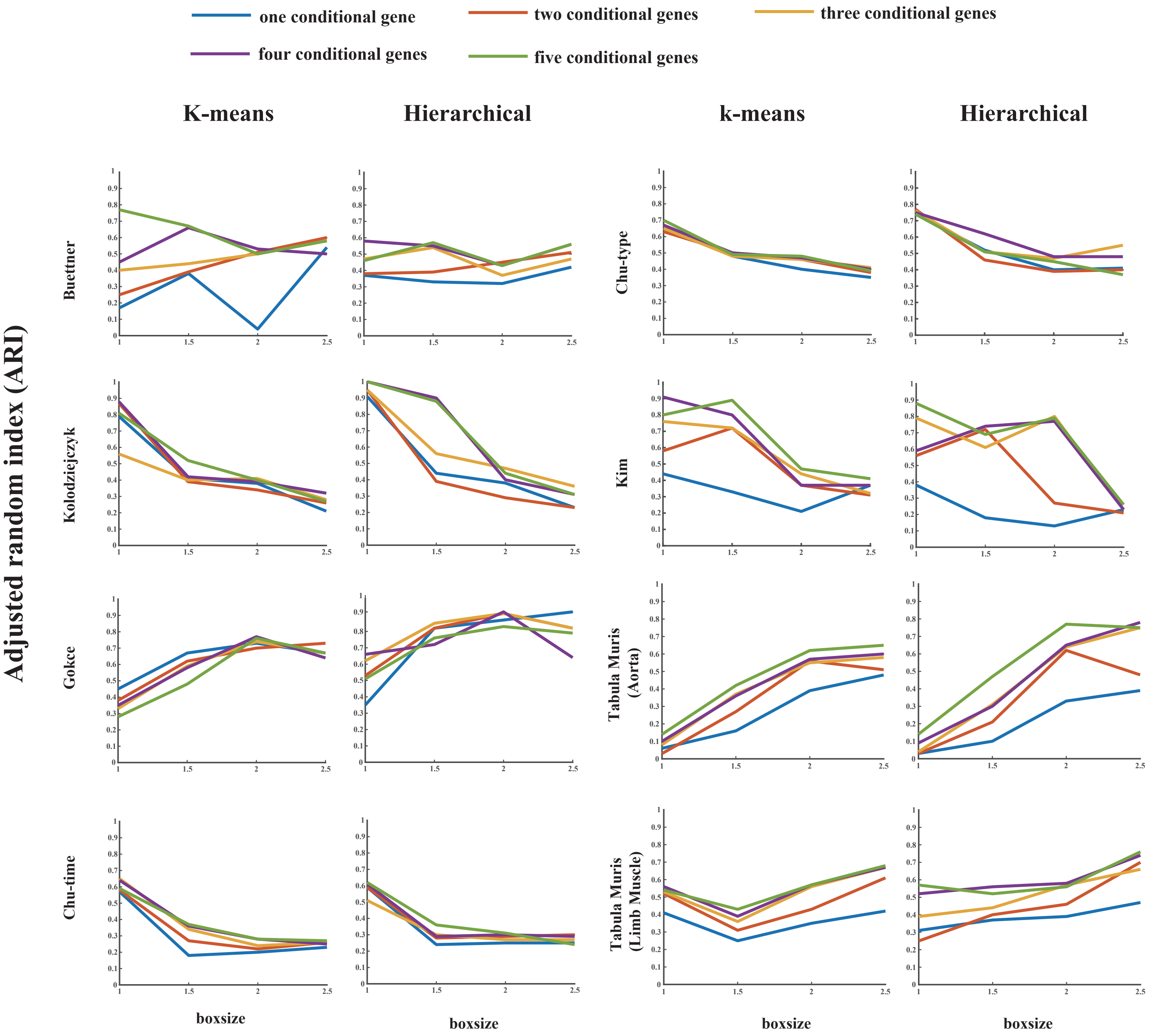

### Figure S8

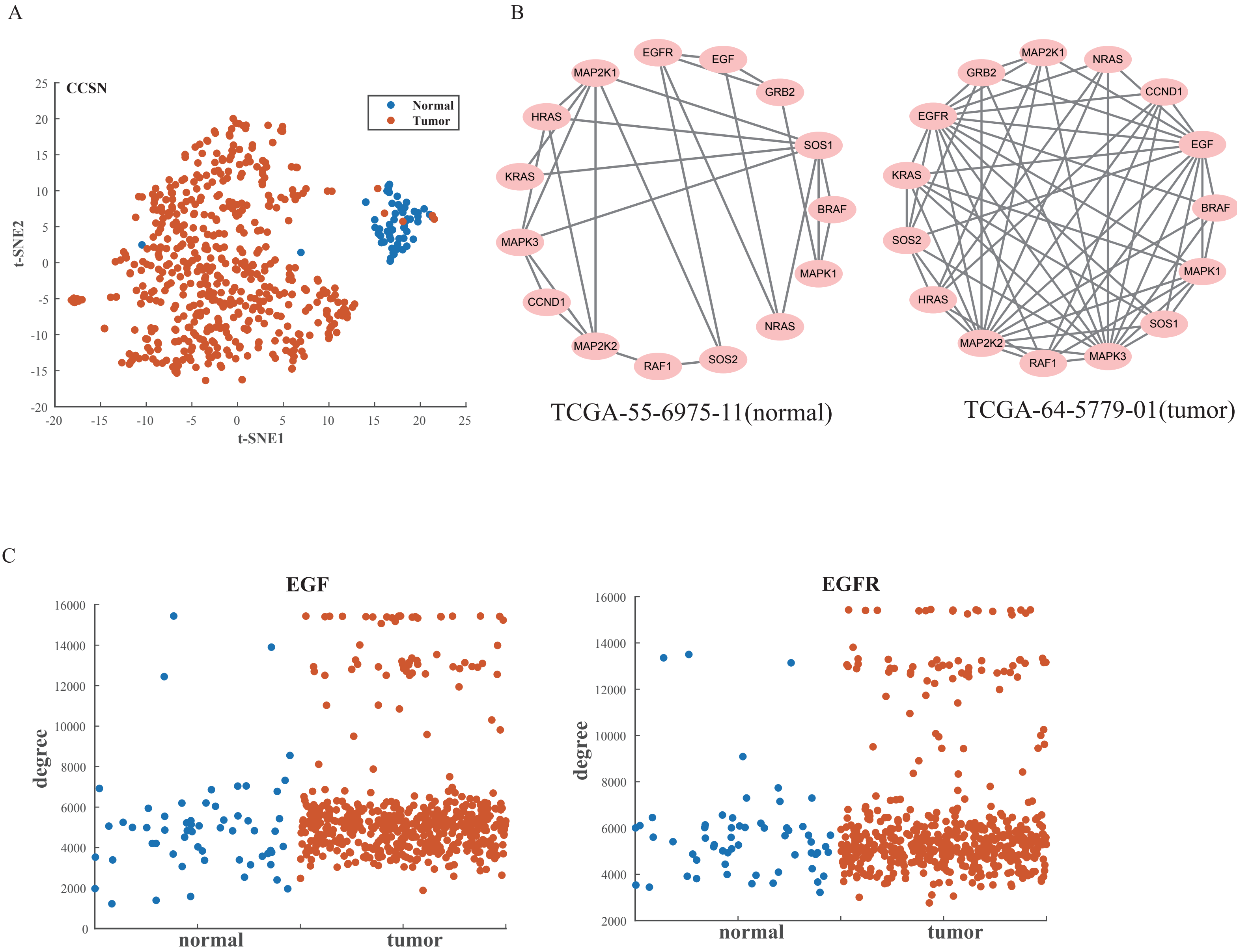
