## Supplementary material for "CCSN: Single Cell RNA Sequencing Data Analysis by Conditional Cell-specific Network": Table S1

**Table S1 The running time of CCSN with different numbers of conditional genes**

|  | **Buettner** | **Yang** | **Chu-time** | **Chu-type** | **10x PBMC** |
| --- | --- | --- | --- | --- | --- |
| N cells | 182 | 447 | 758 | 1,018 | 11,528 |
| 1 conditional gene (h) | 0.33 | 0.38 | 1.75 | 4.25 | 53.9 |
| 2 conditional genes (h) | 0.43 | 1.26 | 2.48 | 5.75 | 53.89 |
| 3 conditional genes (h) | 0.62 | 1.53 | 4.63 | 7.34 | 79.27 |
